# Disulfide disruption reverses mucus dysfunction in allergic airway disease

**DOI:** 10.1101/768424

**Authors:** Leslie E. Morgan, Siddharth K. Shenoy, Dorota Raclawska, Nkechinyere A. Emezienna, Vanessa L. Richardson, Naoko Hara, Anna Q. Harder, Hassan M. El-Batal, Chelsea M. Magin, Diane E. Grove Villalon, Gregg Duncan, Justin S. Hanes, Jung Soo Suk, David J. Thornton, Fernando Holguin, William J. Janssen, William R. Thelin, Christopher M. Evans

## Abstract

Airway mucus is essential for healthy lung defense^1^, but excessive mucus in asthma obstructs airflow, leading to severe and potentially fatal outcomes^2–5^. Current asthma therapies reduce allergic inflammation and relax airway smooth muscle, but treatments are often inadequate due to their minimal effects on mucus obstruction^6,7^. The lack of efficacious mucus-targeted treatments stems from a poor understanding of healthy mucus function and pathological mucus dysfunction at a molecular level. The chief macromolecules in mucus, polymeric mucins, are massive glycoproteins whose sizes and biophysical properties are dictated in part by covalent disulfide bonds that link mucin molecules into assemblies of 10 or more subunits^8^. Once secreted, mucin glycopolymers can aggregate to form plugs that block airflow. Here we show that reducing mucin disulfide bonds depolymerizes mucus in human asthma and reverses pathological effects of mucus hypersecretion in a mouse allergic asthma model. In mice challenged with a fungal allergen, inhaled mucolytic treatment acutely loosened mucus mesh, enhanced mucociliary clearance (MCC), and abolished airway hyperreactivity (AHR) to the bronchoprovocative agent methacholine. AHR reversal was directly related to reduced mucus plugging. Furthermore, protection in mucolytic treated mice was identical to prevention observed in mice lacking Muc5ac, the polymeric mucin required for allergic AHR in murine models^9^. These findings establish grounds for developing novel fast-acting agents to treat mucus hypersecretion in asthma^10,11^. Efficacious mucolytic therapies could be used to directly improve airflow, help resolve inflammation, and enhance the effects of inhaled treatments for asthma and other respiratory conditions^11,12^.

---

With daily exposures to >8,000 liters of air containing billions of particles and potential pathogens, respiratory tissues embody the need for robust host defense. Airway mucus is critical for protection, but poor control of mucus function is central to numerous lung diseases. In patients who die during asthma exacerbations, mucus obstruction is a feature long-recognized by pathologists^13^, with plugging observed in >90% of cases and thus considered a major cause of fatal obstruction^5^. Mucus obstruction is also prominent in non-fatal cases of severe asthma^2^, but effective mucolytic therapies are lacking. Accordingly, asthma treatments could be significantly improved by determining molecular mechanisms of mucus dysfunction^10^.

The predominant macromolecules in mucus are polymeric mucin glycoproteins^10,11^. In health, effective host defense requires homeostatic mucin synthesis and secretion^1^. By contrast, excessive mucin production and secretion are demonstrated by alcian blue-periodic acid Schiff’s (AB-PAS) staining within airway surface and submucosal gland epithelial cells, and in plugs filling small airways in fatal asthma **(Fig. 1a)**. Mucin glycoprotein overproduction is also prominent in mild-to-moderate disease^14^, suggesting a potentially broad etiological role for mucus hypersecretion in asthma.

**Fig. 1.**
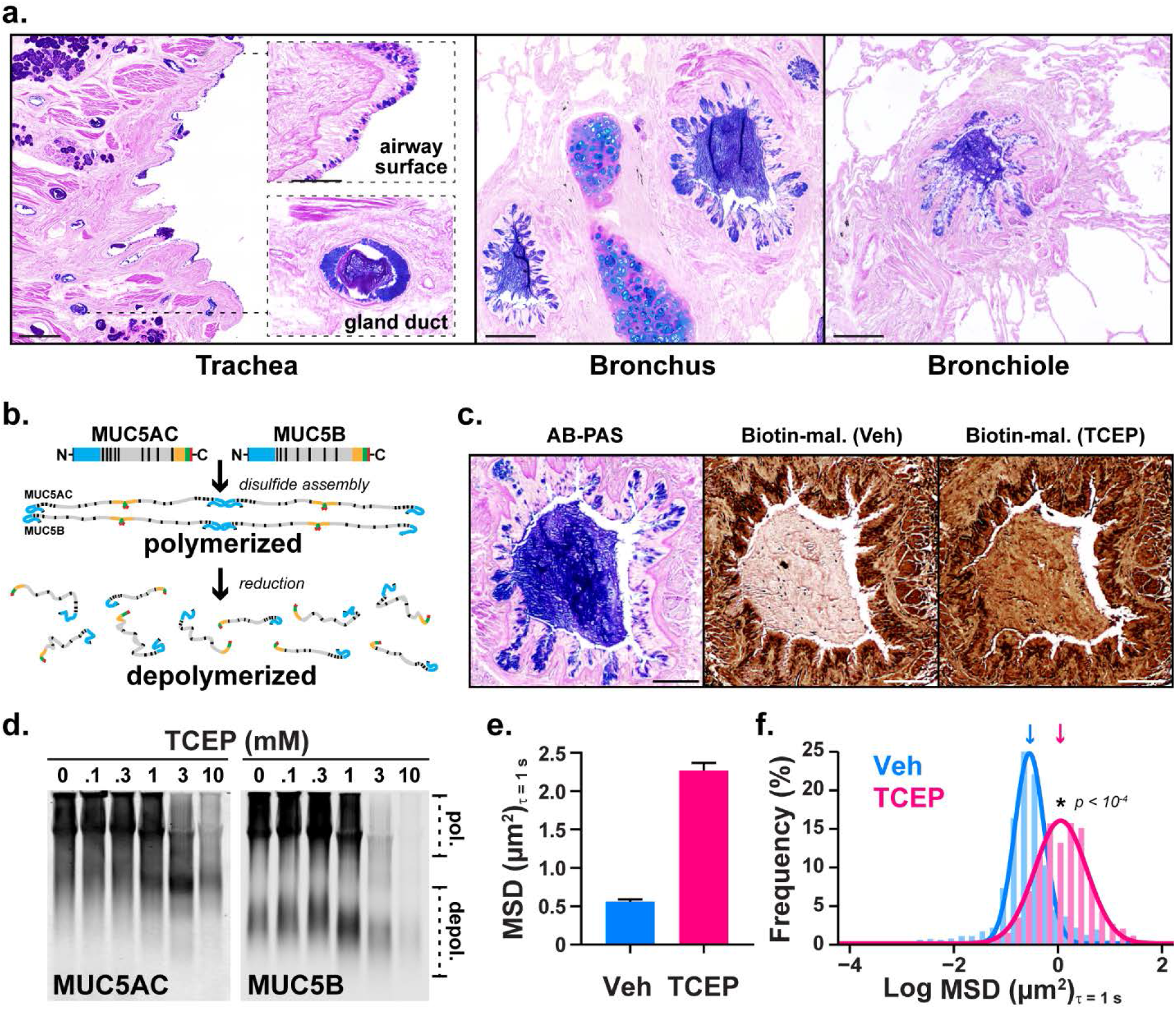
Polymeric mucins in asthmatic airways are targets for disulfide disruption. **a.** AB-PAS stained tissues from the lungs of patients who died during asthma exacerbations demonstrate mucin glycoproteins in large and small airways. Scale bars, 500 μm and 100 μm in insets (trachea), 200 μm (bronchus and bronchiole). **b.** MUC5AC and MUC5B assemble via amino (N-) and carboxyl (C-) terminal disulfide bonds that are sensitive to reducing agents. **c.** A mucus plug in a fatal asthma airway was examined in consecutive sections stained with AB-PAS (purple), or labeled with biotinylated-maleimide (brown) after incubation at 37° C for 10 min with saline vehicle (Veh) or tris(2-carboxyethyl)phosphine (TCEP, 10 mM). Scale bars, 250 μm. **d-f.** Expectorated sputum was treated with TCEP (0.1-10 mM, 37° C, 30 min), separated by electrophoresis (1% SDS/agarose, non-reducing), and detected by immunoblot for MUC5AC and MUC5B (**d**). Note that full reduction causes epitope loss for both anti-mucin antibodies resulting in decreased signal intensities at higher TCEP concentrations in **d**. Diffusion of 2-μm carboxylated micro-particles was evaluated in mucus samples aspirated from fatal asthma bronchi (**e,f**). Compared to vehicle controls (Veh, cyan), particle mean square displacement (MSD) increased significantly after TCEP (magenta). Data in **e** are means ± sem of summarized linear-scale values. Data in **f** are log distributions of particles (n = 1,303 Veh, 1,425 Tx) with curves showing Gaussian non-linear fits (r^2^ = 0.99 for Veh and TCEP). Significance was determined by two-tailed Mann-Whitney U tests with p-value and ‘*’ denoting significance from Veh in **f**. Inverted arrows in **f** indicate locations of median values.

Two polymeric mucin glycoproteins are abundant in airway mucus. The chief airway mucin in healthy mucus is MUC5B, whose absence in mice results in impaired mucociliary clearance (MCC), pathogen accumulation, and spontaneous lethal infections^1^. In humans with asthma, MUC5B decreases in many patients^14,15^, and lower MUC5B levels are associated with worsened disease^15^. On the other hand, excessive MUC5B is also a risk factor for developing pulmonary fibrosis^16,17^, and *Muc5b* has a gene dosage effect on lung fibrosis in mice^18^. The other airway polymeric mucin, MUC5AC, is expressed at low levels at baseline, but it is dramatically up-regulated in human asthma^14,15^ and in mouse models^19^ where it is required for asthma-like mucus obstruction and airway hyperreactivity (AHR)^9^.

Taken together, MUC5AC and MUC5B play significant functions, but their precise roles in health and disease are complex^10^. The genetic, developmental, and environmental factors that cause aberrant *MUC5AC* and *MUC5B* gene expression are active areas of investigation. However, given the inherent heterogeneity among these pathways, finding selective gene expression or signal transduction targets that can prevent mucus dysfunction while still preserving (or improving) mucus defense remains an on-going challenge^10,11,20^. As an alternative, we propose that pathological effects of mucus hypersecretion can be reversed while healthy functions are enhanced by directly targeting mucin glycoprotein polymers.

Polymeric mucins are evolutionary precursors of the hemostasis protein von Willebrand factor (vWF). Accordingly, they possess vWF-like amino (N-) and carboxyl (C-) terminal cysteine-rich domains that form covalent disulfide intermolecular linkages to form large glycopolymers^21^. MUC5AC and MUC5B first assemble in the endoplasmic reticulum as C-terminal disulfide dimers. Then in the Golgi, along with becoming heavily glycosylated, they further multimerize via N-terminal disulfide linkages^8^. Upon secretion, MUC5AC and MUC5B become hydrated, extend into strands, and form a porous mesh-like gel that traps particles and mediates MCC.

Due to the nature of mucus as a gel polymer, it is exquisitely sensitive to changes in the concentrations of solid materials within its matrix. Accordingly, when mucins are overproduced and hypersecreted in asthma, MCC dysfunction reflects aberrant gel behavior that can be corrected by disrupting its polymeric mucin components^11^. Therefore, we tested whether aberrant mucus gel structure, MCC, and airway obstruction in asthma can be improved by disrupting mucin disulfides with the fast-acting reducing agent tris(2-carboxyethyl)phosphine (TCEP) **(Fig. 1b)**.

Using mucus from human asthma patients, we verified the presence of targets that could be reduced with TCEP under physiologic conditions (pH 7.4, 37° C). In histologic samples from fatal asthma, AB-PAS positive plugs were sensitive to TCEP as demonstrated by alkylation of reduced thiols with biotinylated maleimide **(Fig. 1c)**. Furthermore, in mucus aspirated from bronchial airways of the same patient, TCEP reduced the sizes of massive MUC5AC and MUC5B polymers in a concentration dependent manner **(Fig. 1d)**. Based on these findings, we tested whether mucin polymer disruption could improve the physical properties of asthmatic mucus.

We used multiple particle tracking (MPT) to assess mucus biophysical properties by quantifying mean square displacement (MSD) of 2 μm diameter carboxylated micro-particles. In fatal asthma mucus, MSD was significantly increased in samples treated with TCEP **(Fig 1e,f)**. When converted to rheologic parameters^22,23^, these results demonstrated that TCEP treatment rendered mucus less viscoelastic **(Suppl. Data Fig 1a)**, an effect that was driven by a significant reduction in its elastic modulus **(Suppl. Data Fig 1b)**. The rapid depolymerization of mucins and improvement of rheologic properties in asthmatic mucus *in vitro* suggested that TCEP could improve mucus functions in animal model of asthma *in vivo*.

To produce allergic asthma-like inflammatory, mucous, and AHR phenotypes, BALB/c mice were exposed to a fungal allergen, *Aspergillus oryzae* extract (AOE), by aerosol weekly for a total of four challenges^9^. Endpoints were studied 48 h after the last AOE challenge, a time point of robust inflammation and mucin overproduction **(Suppl. Data Fig. 2a)**^9^. Nebulized TCEP was used to determine the effects of mucolytic treatment on mucus properties and functions *in vivo*.

**Fig. 2.**
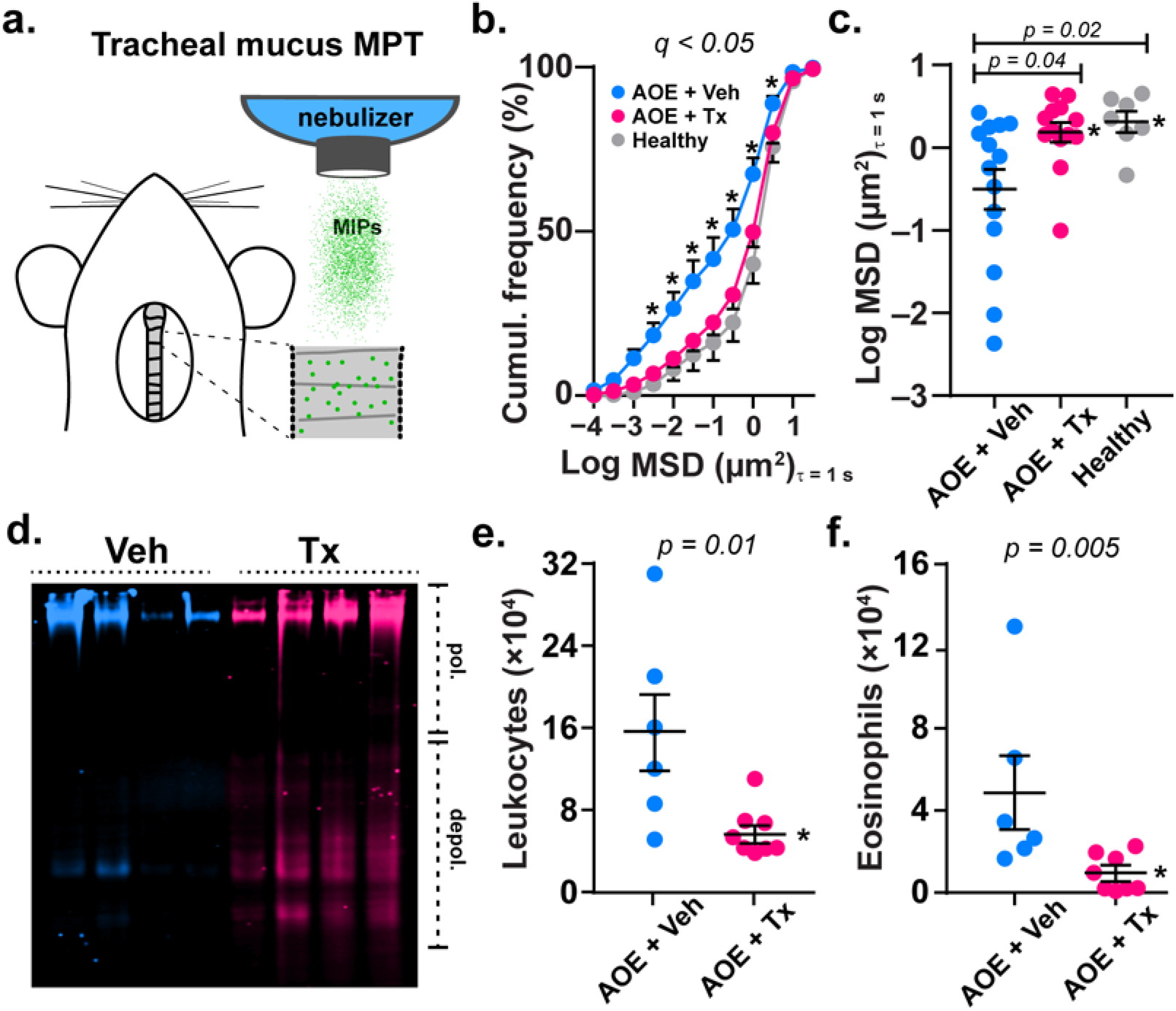
Mucolytic treatment improves mucus function in allergic mouse airways. **a.** Mucus was probed in tracheal preparations *ex vivo* using 100 nm muco-inert particles (MIPs). **b,c.** In AOE-challenged mice receiving 500 mM aerosol TCEP treatment (Tx, magenta, n = 13), MIP diffusion measured as mean square displacement (MSD) increased significantly compared vehicle (Veh, cyan, n = 14) and non-AOE exposed animals (Healthy, gray, n = 7). Cumulative distribution data in **b** show the distributions of MSD values (means ± sem). Scatter plot data in **c** show individual median MSD values per animal. **d-f.** MCC was tested in AOE challenged mice treated by nose-only aerosol with TCEP (Tx, 500 mM) or Veh (cyan, n = 6), followed by immediate lung lavage. Lectin blot analysis using UEA-1 (α1,2-fucose) in **c** shows disruption of mucin polymers in TCEP-treated animals (magenta) compared to vehicle (cyan). Image shows 4 samples per group. Numbers of total leukocytes (**d**)and eosinophils (**e**)recovered in lung lavage decreased significantly in Tx (magenta, n = 8) vs Veh (cyan circles, n = 6) exposed animals. Lines and error bars in **c,e,f** are means ± sem. ‘*’ denotes significance using a cut-off of 0.05 determined by unpaired t-tests using a two-stage step-up method at a 5% false discovery rate from Veh in **b**, and by two-tailed Mann-Whitney U test in **c,e,f**.

To evaluate mucus directly on mouse airway surfaces, we employed MPT to quantify MSD of muco-inert nano-particles (MIPs) aerosolized onto mouse tracheas *ex vivo*^24–26^. Tracheas were removed from saline or AOE challenged mice, opened, placed on glass coverslips, and treated with nebulized 100 nm diameter MIPs suspended in saline vehicle or TCEP **(Fig 2a)**. Compared to uninflamed controls, MIP diffusion was impaired in the tracheal mucus of AOE-challenged mice **(Fig. 2b)**, resulting in a 2.7-fold decrease (p = 0.02) in median MSD values **(Fig. 2c)**. Upon mucolytic treatment, MSD normalized to levels indistinguishable from non-allergic controls **(Fig. 2b,c)**, reflecting an increase in computed mucus mesh spacing **(Suppl. Data Fig. 3)**. Taken together, these findings suggested that mucolytic treatment normalized mucus gel microstructure, which we postulated would also improve mucociliary function *in vivo*.

**Fig. 3.**
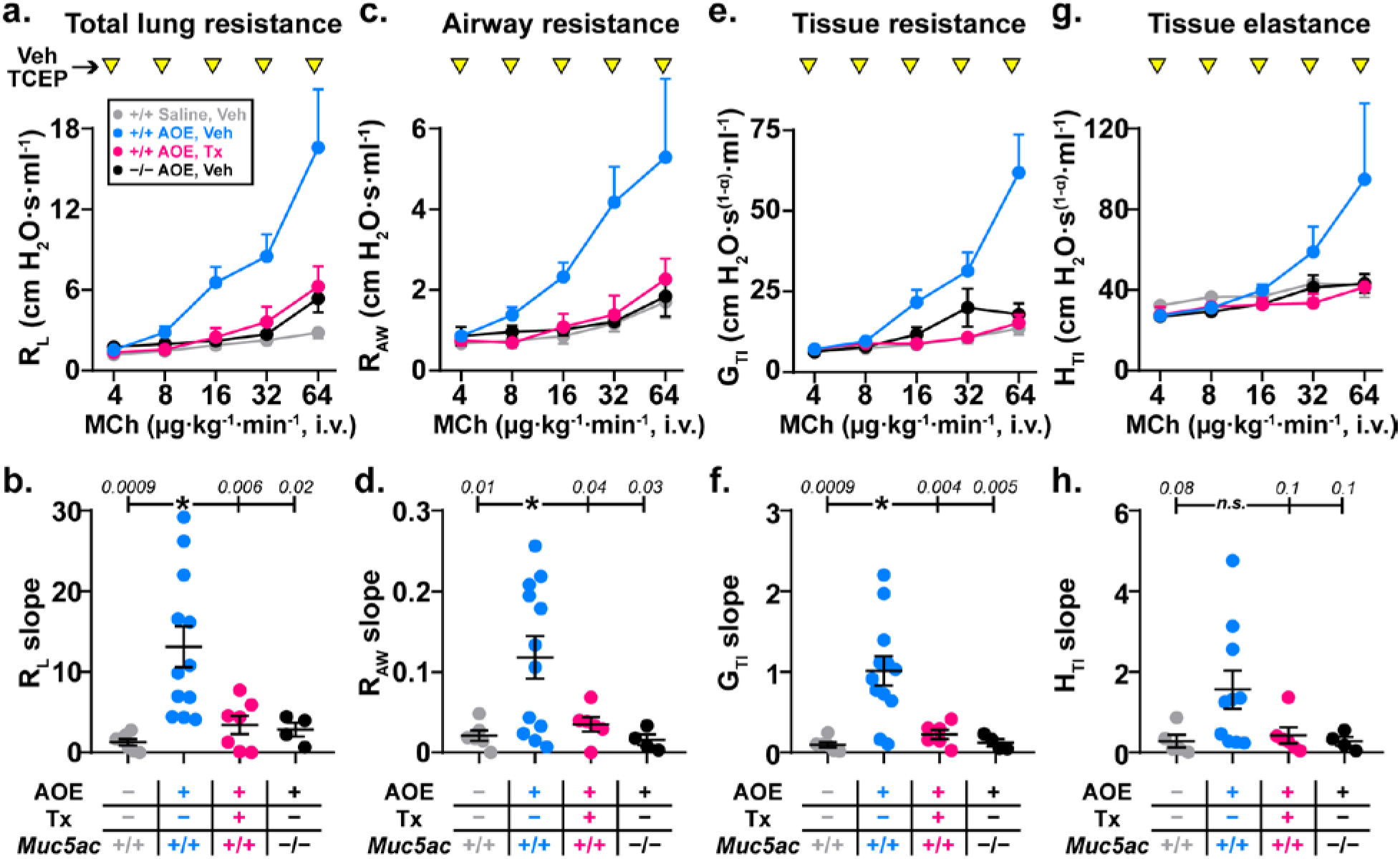
Mucolytic treatment reverses allergic airway hyperreactivity. Dose response curves to MCh (4-64 μg/kg/min, i.v.) were generated in AOE challenged allergic WT mice (magenta, n = 7) treated with TCEP between MCh doses (100 mM, inverted yellow triangles). For comparison, saline challenged non-allergic WT mice (gray, n = 6), AOE challenged allergic WT mice (cyan, n = 12), and AOE challenged *Muc5ac*^−/−^ mice (black, n = 4) were treated with nebulized vehicle (Veh) during i.v. MCh dose response tests. For each mouse, dose response curves were fitted by log-linear best-fit regression analysis, and slopes of regression lines were analyzed by one-way ANOVA. ‘*’, p < 0.05 using Dunnett’s test for multiple comparisons relative to AOE-challenged Veh-treated WT mice (p-values are shown). Total lung resistance (R_L_ in **a,b**), conducting airway resistance (R_AW_ in **c,d**), tissue resistance (G_TI_ in **e,f**) and tissue elastance (H_TI_ in **g,h**) are shown.

We thus investigated the efficacy of inhaled mucolytic treatment on MCC in allergically inflamed mice *in vivo*. Animals were allergen challenged with AOE, and 48 h after the challenge, they received TCEP via nose-only aerosol (see **Suppl. Data Fig. 2b**). Immediately after mucolytic treatment, lungs were lavaged, and the disruption of mucin polymers and elimination of inflammatory cells from airspaces were assessed^18^. In TCEP-treated mice, mucins demonstrated faster electrophoretic migration relative to controls thereby validating effective depolymerization **(Fig. 2d** and **Suppl. Data Fig. 4)**. Concordant with mucolysis, there were acute decreases in inflammatory cell numbers in lung lavage fluid **(Fig. 2e,f** and **Suppl. Data Table 1)**. The rapid elimination of leukocytes from airway surfaces upon mucolytic treatment provided direct functional evidence of improved MCC^18^.

**Fig. 4.**
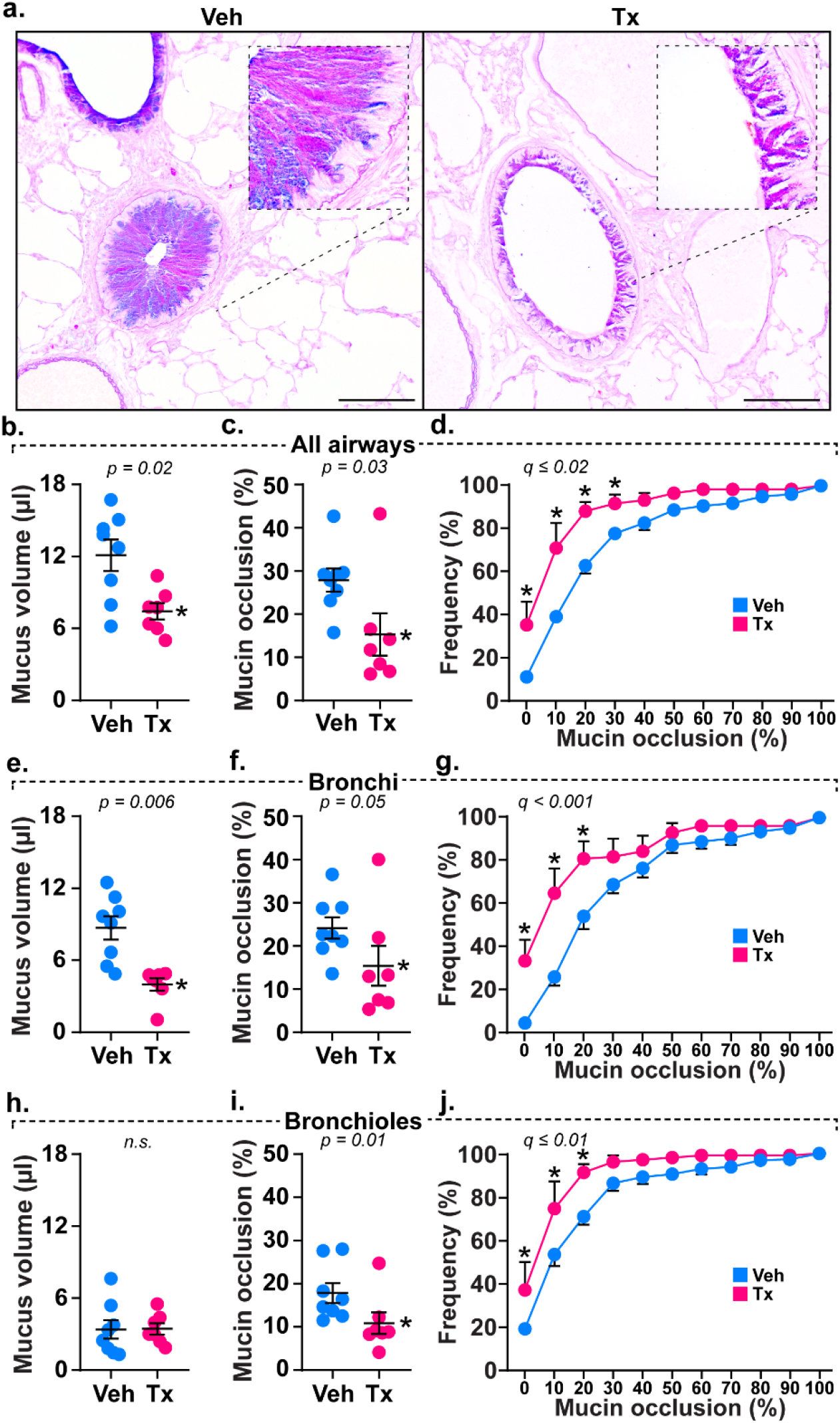
Mucolytic treatment disrupts mucus plugging. **a.** AB-PAS stained lungs obtained after MCh dose response tests show mucin glycoproteins obstructing airspaces in vehicle (Veh) treated mice that are disrupted by mucolytic treatment (Tx). Scale bars, 500 μm (low power) and 25 μm (high power). **b-j.** Calculated mucus volumes (**b,e,h**), mean fractional occlusion (**c,f,i**), and heterogeneous plugging (**d,g,j**) were significantly decreased in AOE-challenged mice receiving mucolytic (Tx, magenta, n = 7) compared to controls (Veh, cyan, n = 8). Mucolytic treatment significantly reduced obstruction across all airways (**b-d**), and this effect was most prevalent in bronchi (**e-g**). Although total mucus volume in bronchioles was low (**h**), occasional mucus aggregates were obstructive and are sensitive to TCEP treatment (**i,j**). Data in **b,c,e,f,h,** and **i**are means ± sem on scatter plots, with p-values shown and significance (*) indicating p < 0.05 by two-tailed Mann-Whitney U test. Cumulative frequency distributions in **d, g,** and **h** show percentages of airways demonstrating occlusion, with ‘*’ demonstrating significance by t-test using a two-stage step-up method at a 5% false discovery rate.

Collectively, our *in vitro*, *ex vivo*, and *in vivo* results showed that mucolytic treatment reversed mucus dysfunction in a setting of allergic inflammation and excessive mucin production. These findings led us hypothesize that mucolytic treatment could also protect against asthma-like airflow obstruction. To test this, we developed a two-route challenge and treatment analysis, with MCh administered intravenously (i.v.) to cause obstruction, and with TCEP administered by inhalation to cause mucolysis. MCh and TCEP treatments were given during pulmonary function tests **(Suppl. Data Fig. 2c)**.

Since we previously found that Muc5ac absence prevents AHR to inhaled MCh, which results in both airway smooth muscle contraction and mucin secretion^9^, we first tested whether Muc5ac was similarly required for AHR to MCh administered intravenously (without mucolytic treatment). Compared to non-allergic animals, AOE challenged mice demonstrated significantly exaggerated increases in total lung resistance (R_L_) and airway resistance (R_AW_) in response to i.v. MCh **(Fig. 3a-d)**. Importantly, these AHR responses were abolished in *Muc5ac* gene deficient animals **(Fig. 3a-d)**, thereby validating a role for mucus hypersecretion in this MCh challenge model. The degrees of protection observed in this chronic mucus prevention setting also established benchmarks for comparing effects of acute mucolytic rescue. Accordingly, we next tested whether inhaled TCEP protected mice from AHR by reversing mucus plugging.

In AOE exposed wild type mice treated with inhaled TCEP during i.v. MCh challenge, we observed significant improvements in R_L_ and R_AW_ **(Fig. 3a-d)**. There was also significant protection of resistance (G_TI_) and elastance (H_TI_) in peripheral lung tissues **(Fig. 3e-h)**. Thus, inhaled TCEP was not acutely detrimental to pulmonary surfactant function, and it appeared to confer protection from AHR by preserving airway patency. Indeed, the protective effects of TCEP treatment in allergic wild type mice were indistinguishable from benchmarks in allergic *Muc5ac*^−/−^ mice (p = 0.98), strongly supporting the hypothesis that mucolytic protection from AHR was linked to reversal of mucus plugging.

To validate the role of mucolysis in protection from AHR in TCEP-treated mice, we confirmed mucin polymer reduction by examining lung lavage fluid via immunoblot **(Suppl. Data Fig. 5)**. We also quantified mucin accumulation in airspaces histologically^9^. During MCh-induced bronchoconstriction, secreted mucin volume, airway obstruction, and heterogeneous plugging were significantly reversed throughout the airways of TCEP-treated animals **(Fig. 4)**. Thus, protection from AHR was directly related to the ability of mucolytic treatment to reduce airway plugging.

In aggregate, the studies reported here show that disrupting mucin polymers improves mucus microstructure, enhances mucus transport, and protects airflow in allergic asthma settings. In patients, mucin overproduction and pathologic changes in mucus biophysical properties are correlated with asthma exacerbations and fatalities^2,5,10,14,27^. These features are observed frequently by pathologists, but they are usually ignored during clinical assessments since diagnostic tests and interventions have lagged. On the diagnostic side, mucus obstruction is becoming more recognizable through high resolution imaging^2^, but treatment options are still constrained a lack of efficacious mucolytics.

Indeed, while potentially beneficial in laboratory settings, mucolytics and expectorants are not widely used therapeutically. The only FDA-approved reducing agent available as an inhaled mucolytic is N-acetylcysteine (NAC), but efficacy is low due to NAC’s weak activity at airway pH and high mucin concentrations^28–30^. We propose that inhaled mucolytic agents can reduce plugging and thus improve airway function in asthma. Nonetheless, it will be critical for any strategy that reverses the detrimental effects of mucus hypersecretion to do so while also protecting mucus-mediated defense. Since disulfide assembly is a critical process in the formation of viscoelastic mucus, reduction remains an attractive target for disrupting mucus plugs in human airways (see **Fig. 1**).

In contrast to the findings reported here, a recent study investigating mucociliary transport showed inhibitory effects of TCEP on mucus transport in non-diseased pigs^31^. However, that investigation focused on tracheobronchial glandular secretions, and it was conducted using large (500-600 μm diameter) metal particles. Given the loads imparted by these materials and the forces required to displace them, inhibitory consequences of TCEP treatment on exogenous transport in that report may not be directly comparable to findings investigating endogenous clearance here. In our studies, TCEP treatment normalized mucus, improved MCC, and reversed acute airway plugging (see **Figs. 2-4**). Thus, even under diseased conditions, mucolytic effects were examined within physiologic constraints.

Our findings suggest that an inhaled mucolytic treatment could confer acute protection from obstruction in asthma. Airway narrowing initiated by smooth muscle contraction is clearly important, and obstruction is amplified by mucus^9^. In bronchoalveolar lavage (BAL) fluid from patients with mild-to-moderate asthma, TCEP rapidly depolymerizes MUC5AC and MUC5B (**Suppl. Data Fig. 6**), demonstrating the presence of targets in asthma patients even under stable disease conditions.

In addition to disulfide targets formed during mucin biosynthesis, there are other potential targets for mucolytic intervention. These include non-disulfide covalent linkages such as sugars added during glycosylation, as well as non-covalent mucin polymer interactions formed during mucin packaging into secretory granules. In addition, in the post-secretory environment oxidant-mediated mucin cross-linking is observed in cystic fibrosis (CF)^32^ and asthma^2^, and it is reported to occur on free thiols in these in static mucus aggregates in these settings. Thus, a disulfide reduction strategy similar to what was applied here for acutely secreted mucus could also improve airway functions under conditions of chronic mucus accumulation^12^. Additional non-covalent mucus interactions associated with macromolecules such as DNA^33^, or effects mediated by ionic and pH environments could be also targeted^34–36^. Although these are not directly affected by reduction *per se*, agents that disassemble disulfides could be applied in combination with other therapies to improve mucus hydration and transport. Thus, treatments that reverse mucus dysfunction could then be considered as possible adjunct or stand-alone therapies.

Having identified the ability to deliver a mucolytic agent, promote MCC, and reverse AHR in a mouse model of allergic asthma, findings reported here could be extended to other lung pathologies where mucus dysfunction is significant^10,11^. Inflammatory or injurious causes of mucus dysfunction vary within and across diseases, yet they still result in a similar outcome—excessive polymeric mucin production. Indeed, mucus allergic asthma exhibited comparable microstructure compared to CF sputum, and both samples had almost identical concentrations of mucus solids (**Suppl. Data Fig. 7**). To this end, even in diseases with diverse primary causes, there could be benefits that derive from optimizing airway mucus. For example, in addition to reducing obstruction, mucin depolymerization could enhance the deposition and absorption of inhaled bronchodilator, anti-inflammatory, or antibiotic agents. Further studies focused on developing effective mucolytic therapies and potentially mucin-selective treatments are needed.

## Materials and Methods

### Human Specimens

BAL fluid was obtained from volunteers with asthma recruited for a University of Colorado Institutional Review Board (IRB) approved study (ages 18-53, n = 2 male, 2 female). BAL was collected from the left upper lobe (lingula projection), and a 2 ml sample was separated and stored “neat” (without centrifugation) to preserve high molecular weight mucins that sediment at low centrifugation speeds.

Mucus aspirated from the lobar bronchi of a patient who died during a fatal asthma exacerbation was obtained from lungs donated to National Jewish Health. Spontaneously expectorated CF sputum samples were collected from patients at the adult CF clinic at Johns Hopkins University. Studies were approved by the Johns Hopkins IRB, and patients gave informed consent to participate.

Tissue sections were made from samples of fatal asthma lung and airway tissues obtained at National Jewish Health from the fatal asthma patient above, and from a second patient whose lungs were donated at Oregon Health and Sciences University (kindly provided by Dr. David Jacoby). Paraffin sections were used for mucin labeling as described below.

### Mucin Protein Detection

Histochemical staining with alcian blue-periodic acid Schiff’s (AB-PAS) stain was performed using standard techniques, as described previously^37^. For immuno-detection, human and mouse mucins were detected using rabbit-anti-human MUC5AC (MAN5AC)^38^, mouse-anti-MUC5AC (45M1, ThermoFisher), rabbit-anti-human MUC5B (H300, Santa Cruz), rabbit-anti-mouse Muc5ac (UNC294)^39^, and rabbit-anti-mouse Muc5b^1^. For biochemical labeling, mucins were also detected using biotinylated *Ulex europaeus* agglutinin I (UEA-I) lectin to detect fucose residues (Vector, Burlingame, CA) or with biotinylated maleimide to alkylate and detect reduced sulfhydryls (ThermoFisher).

### Immuno-/ Lectin Blotting

Lung lavage and mucus specimens were separated on SDS/agarose gels to determine mucin profiles and changes in mucin polymer sizes upon reduction^40^. A dot-blot ELISA was performed to assess MUC5B levels in each human lavage sample, and lanes were equilibrated to contents of MUC5B, the predominant mucin in these patients with non-exacerbated asthma. In allergic mice, although lung lavage fluid contains mixtures of both Muc5ac and Muc5b, they are found at higher concentrations and can be more homogeneously sampled (see below). Therefore, mouse lung lavage samples were loaded into gels using equal volumes. After electrophoresis and transfer to PVDF membranes by vacuum blot^40^, mucins were probed with anti-mucin antibodies (1:1,000-5,000 dilutions as indicated), and then detected with goat anti-mouse IRDye 680RD or goat anti-rabbit IRDye 800CW antibodies (LI-COR, Lincoln, NE). Image analysis was performed using Image Studio software (LI-COR).

### Particle Tracking in Human Mucus

For human airway mucus samples, 2.0 μm fluorescent carboxylated microspheres (ThermoFisher) were used to estimate mucus microrheology. Frozen mucus samples from the patients above were thawed on ice, distributed into 100 μl aliquots. A 10 μl solution of TCEP (100 mM) or saline (vehicle) containing 100,000 microspheres was added to each mucus sample. Samples were incubated at 37° C for 10 min.

Immediately after incubation, samples were loaded onto chambered slides and video imaged at 5 randomly chosen sites on a BX63 microscope for 30 sec (7.5 frames per sec) using a DP80 camera (Olympus, Center Valley, PA). Particle tracking was performed using Olympus cellSens software, and analyses were made to derive MSD values using a custom Matlab program (Mathworks, Natick, MA). Using the frequency-dependent Stokes-Einstein equation, we translated MSD values to obtain approximations of micro-rheological properties of airway mucus based on previously published methods^22,23^.

### Mice

Studies were conducted with approval of the University of Colorado Denver and Johns Hopkins University Institutional Animal Care and Use Committees. Male and female BALB/cJ wild type mice were purchased from the Jackson Labs (Bar Harbor, ME). *Muc5ac*^−/−^ mice were previously crossed onto a congenic BALB/cJ strain background^9^. Animals were housed under specific pathogen-free conditions and used in allergic asthma studies beginning at ages 6-8 weeks. To induce allergic inflammation and mucin overproduction, mice were challenged using aerosolized Aspergillus oryzae extract (AOE; Sigma), as reported previously^9^ and shown in **Suppl. Data Fig. 2**. Mice received four weekly challenges, and endpoint analyses were studied 48 h after the last AOE challenge. Mice exposed to saline challenges were used as controls.

### Particle Tracking in Sputum Samples and Freshly Excised Mouse Tracheas

MIPs were prepared using a previously described method^25^. Briefly, 5-kDa methoxy-polyethylene glycol (PEG)-amine (Creative PEGWorks, Chapel Hill, NC) molecules were densely conjugated to the surface of 100 nm carboxyl-functionalized polystyrene beads (PS-COOH; Product#: F8801, 580/605, Molecular Probes, Eugene, OR), by covalent attachment to the carboxyl end groups on the PS-COOH particles using a crosslinker, 1-ethyl-3-(3-dimethylaminopropyl) carbodiimide hydrochloride (Sigma-Aldrich, No. E7750) and an amide, sulfo-N-hydroxysulfosuccinimide sodium salt (Sigma-Aldrich, No. 56485) in borate buffer (200 mM, pH 8.0, Growcells, No. MRGL-1100). The physicochemical properties of MIPs were then measured by a Zetasizer Nano ZS90 (Malvern Analytical, Malvern, UK). The resulting 100 nm MIPs exhibited hydrodynamic diameters of 98.0 ± 6.9 nm, polydispersity indices of 0.1 ± 0.01 and -potentials (i.e. indicative of particle surface charges) of −4.1 ± 0.2 mV measured in 10 mM NaCl pH 7.4 at 25°C.

CF and asthmatic mucus samples were stored at 4°C immediately after collection for up to 24 hours for particle tracking experiments to ensure that the inherent mucus microstructure is preserved^41^. Aliquots of 30 μL of sputum were added to custom microscopy chambers. Next, 0.5 μl of MIP (0.00001% v/v) were mixed gently with sputum sample to evenly distribute particles within the sample. The chamber was sealed with a circular coverslip and incubated at room temperature for 30 min prior to imaging.

Fluorescent MIPs were administered to the mucosal surfaces of tracheas freshly dissected from mice. Briefly, tracheal tissues harvested from animals were cut along the dorsal cranial-caudal axis and laid flat to expose mucosal surfaces. MIPs were diluted in saline (i.e. vehicle control) or TCEP (500 mM) at 0.02% w/v and administered in <1 μl volumes onto mucosal surfaces of the flat-mounted tracheas using a vibrating mesh nebulizer (Aerogen Solo, Chicago, IL) controlled by an Analog Discovery 2 data acquisition device (Digilent, Pullman, WA). After treatment with MIPs in vehicle or TCEP, tracheas were laid face down on a coverslip-bottomed dish (Thermo Scientific) and sealed with vacuum grease and parafilm, followed by incubation for 30 min at 4°C to temporarily immobilize cilia prior to particle tracking experiments^42^.

Motions of MIPs were captured at 15 frames per second (i.e. exposure time of 67 ms) for 20 seconds on an Axiovert D1 inverted epifluorescence microscope (Zeiss, Chicago, IL) equipped with a Photometrics Evolve EMCCD 512 camera (Photometrics, Tucson, AZ) and a MetaMorph software (Molecular Devices, San Jose, CA). MSD values are averages of squared distances traveled by individual particles at a given time interval (i.e. timescale) and thus are directly proportional to particle diffusion rates. Tracking resolution was determined by analyzing MSD on 100 nm MIP particles immobilized in glue between two coverslips, which was found to be less than Log10(MSD_τ = 1s_) of −3.

### Acute Endogenous Clearance and Mucolytic Testing

Acute endogenous clearance was performed as described^18^. In the present studies, AOE challenged mice were exposed to TCEP (500 mM) or saline aerosol for 40 min, at the end of which they were immediately used in lung lavage studies. Animals were anesthetized with urethane (2 g/kg, i.p.) and then tracheostomized with a 20 gauge blunt tip Luer stub^40^. Within 5 min of removal from aerosol treatment, lavage was performed using saline to isolate mucins and inflammatory cells. AEC was determined by quantifying an acute decrease in leukocyte numbers in lavage fluid in TCEP relative to saline treated animals.

Lavage macrophages, lymphocytes, neutrophils, and eosinophils were enumerated as described above for human BAL. Remaining lavage fluid was divided into two aliquots. One portion was centrifuged at 1,200 rpm for 10 min at 4° C for subsequent studies of cytokines and other soluble factors. The other portion was treated with 1 M iodoacetamide (1/100 volume) to quench drug activity and alkylate thiols liberated by disulfide reduction; this portion was stored neat. Both were frozen on dry ice and stored at −80° C.

### Airway Hyperreactivity Measurements

Mice anesthetized with urethane (2 g/kg, i.p.), tracheostomized with a beveled blunt Luer stub, placed on a flexiVent (Scireq, Montreal, Quebec). Ventilated mice were paralyzed with an initial 0.2 ml injection of succinylcholine chloride (300-35 mg/kg, i.p.) to prevent spontaneous respiration. Mice were then cannulated with a 20 gauge catheter in the inferior vena cava. To maintain paralysis throughout the experiment, a continuous infusion of succinylcholine was administered as described below.

Total lung resistance (R_L_), airway resistance (R_AW_), tissue resistance (G_TI_), and tissue elastance (H_TI_) were measured at baseline and in response to methacholine (MCh), which induces bronchoconstriction and mucin secretion^9^. MCh was diluted to a final concentration of 1 μg MCh/g body weight/ml of normal saline solution also containing succinylcholine chloride (20 μg/ml final concentration). This was administered using a digital infusion pump with rates adjusted to 4, 8, 16, 32, and 64 μl per min. MCh doses were given for 4-5 min with recordings of lung function made 20 times during this period. Between doses of MCh, TCEP (500 mM) was administered through an in-line ultrasonic nebulizer (10 s per dose, ~1-2 μl delivery to the lungs per dose). After the last dose of MCh was delivered, lungs were either lavaged with saline to evaluate mucin depolymerization by immunoblot, or fixed to assess mucus plugging^40^.

### Histology

To assess plugging, lungs were fixed transmurally at end-expiratory volume during responses to the highest dose of MCh used. Lungs were fixed in methacarn for 24 h, then transferred to absolute methanol. Fixed lungs were excised after 30-60 minutes and placed in a scintillation vial filled with methacarn. Lung volume was calculated using volume displacement in absolute methanol. Lungs were then cut into 2 mm cubes (~30 per lung), paraffin embedded in random orientations, and sectioned^9,40^. AB-PAS stained tissues were then scanned on an Olympus BX63 microscope, and each tissue fragment was imaged from the center using a 10x objective. Total airway vs. parenchyma volume fractions were determined using a 250 × 250 μm grid, and mucin volume fractions were determined using a 50 × 50 μm grid. Volume fractions were then normalized to lung and airway volumes.

### Statistical Analysis

Statistical analysis and graphs were performed in Prism (GraphPad, San Diego, CA). Comparisons were made using t-Tests, Mann-Whitney U tests, or ANOVA with post-hoc analyses as noted.

## Acknowledgements

This study was funded by NIH grants HL080396 and ES023384 (C.E.), HL130938 (C.E. and W.J.) and by NIH grant HL125169 and Cystic Fibrosis Foundation grant HANES16XX0 (J.H.).

## Author contributions

L.M., S.S., C.M., J.H., J.S., D.T., F.H. W.J. W.T. and C.E. designed the study; L.M., S.S., D.R., N.E., V.R., N.H., A.H., H.E., D.V., and G.D. planned, performed, and analyzed experiments. L.M. and C.E. wrote the manuscript with help from all coauthors. C.E. supervised the manuscript preparation.

## Conflict of Interest Statement

The muco-inert particle technology described in this article is being developed by Kala Pharmaceuticals. J.H. declares a financial, a management/advisor, and a paid consulting relationship with Kala Pharmaceuticals. J.H. is a cofounder of Kala Pharmaceuticals and owns company stock, which is subject to certain restrictions under Johns Hopkins University policy. C.E. is a paid consultant with Eleven P15, a company focused on early detection and treatment of pulmonary fibrosis. The terms of these arrangements are being managed by Johns Hopkins University and the University of Colorado in accordance with respective institutional conflict-of-interest policies. W.T. and D.V. are employees of Parion Sciences, Inc., a company that designs and tests novel mucolytic agents. No proprietary mucolytic agents were used in this study. All other authors declare no conflicts of interest.

## Data availability

The data that support the findings of this study are available from the authors on reasonable request, see author contributions for specific data sets.

---

**Suppl. Data Fig. 1.**
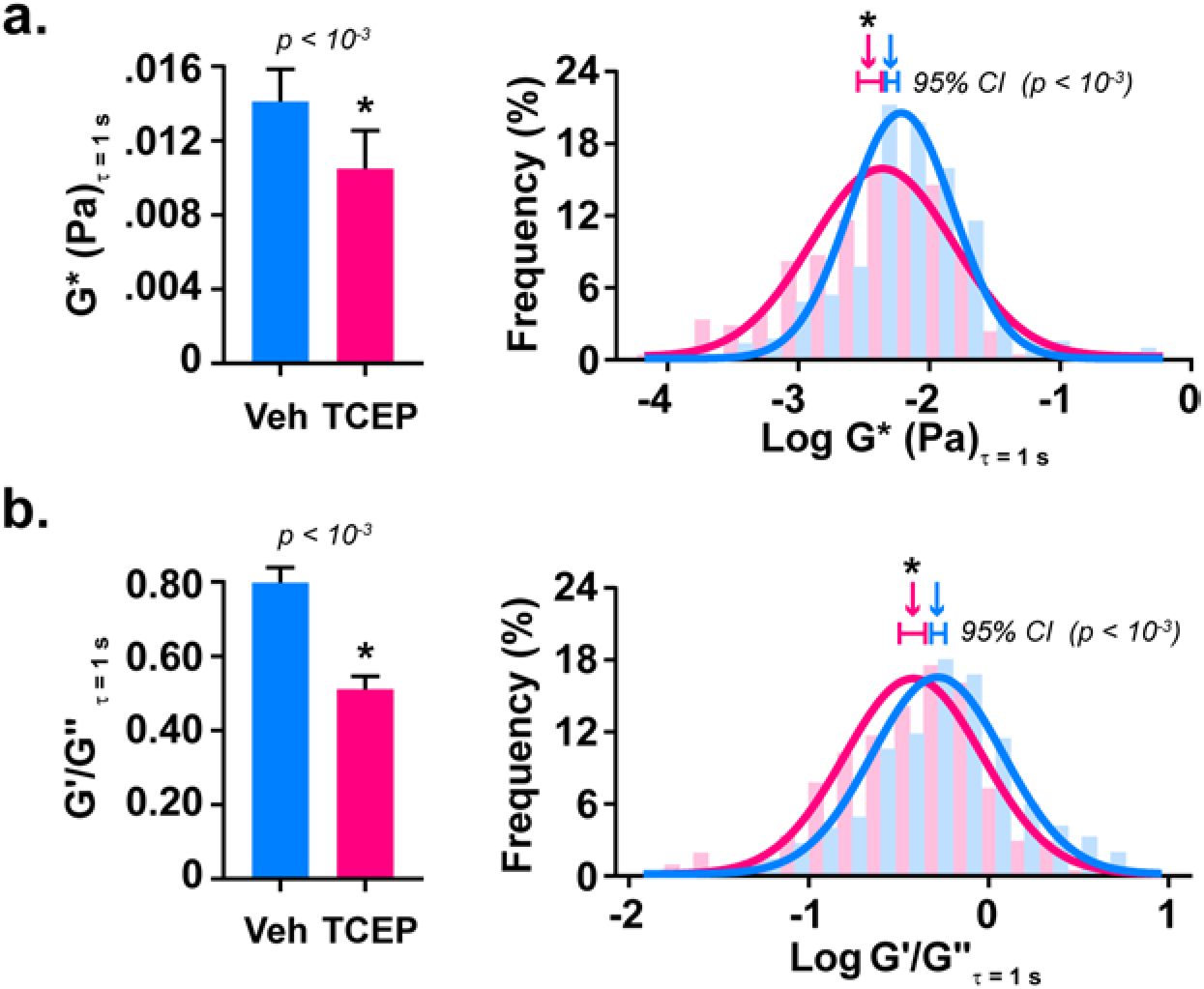
TCEP reduces mucus viscoelasticity. MPT was used to estimate mucus rheology. Carboxylated 2 μm diameter polystyrene micro-particles were applied to mucus and incubated in the presence of TCEP (10 mM, magenta) or saline vehicle (Veh, cyan) for 30 min at 37° C. MSD values were mathematically converted to complex (G*), elastic (G’), and viscous (G’’) moduli. Compared to Veh controls, G* (**a**) and G’/G” ratio (**b**) decreased significantly after TCEP. Bar graphs show means ± sem of summarized linear-scale values. Histograms show log distributions of particles (n = 549 Veh, 205 Tx) with Gaussian fits, with horizontal bars indicating 95% confidence intervals and inverted arrows indicating median values. Significance was determined by two-tailed Mann-Whitney U tests with p-values shown and ‘*’ denoting significance between TCEP and Veh treated samples.

**Suppl. Data Fig. 2.**
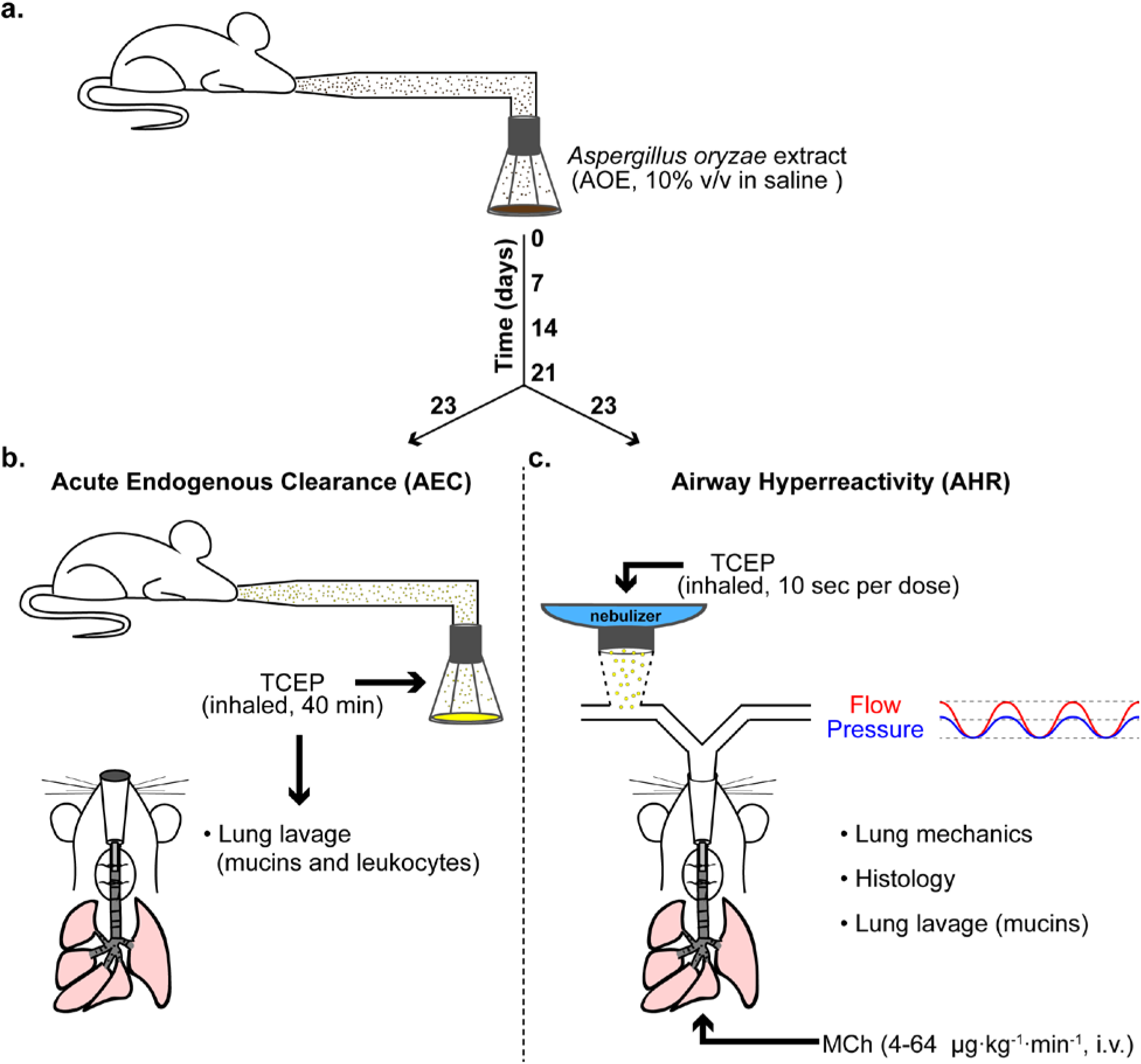
Mouse models used for *in vivo* studies. **a.** Wild type (WT) BALB/cJ mice were challenged by nose-only aerosol with *Aspergillus oryzae* extract (AOE) diluted 1:10 (vol/vol) in saline. A total volume of 5 ml was delivered over a 40 min period on days 0, 7, 14, and 21. Control mice received nose-only aerosols containing saline only. Mice were used for testing acute endogenous clearance (AEC) or airway hyperreactivity (AHR) two days after their last AOE challenge (day 23). **b.** For AEC, TCEP (500 mM) or saline vehicle was delivered in a 5 ml aerosol for 40 min. Mice were immediately used in lung lavage studies, and inflammatory cell numbers were enumerated. **c.** For AHR, lung mechanics and AHR to methacholine (MCh) were assessed in the presence of TCEP (100 mM) or saline vehicle, followed by determination of mucus plugging and mucin disruption.

**Suppl. Data Fig. 3.**
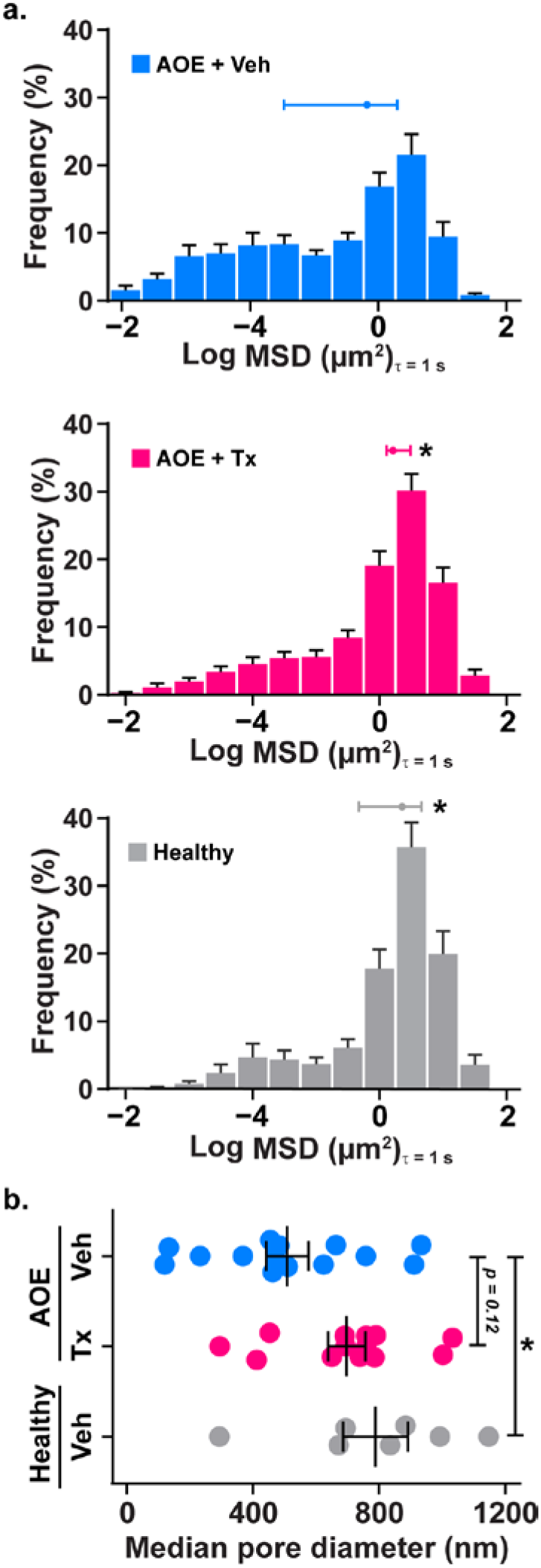
Microstructural properties affected by TCEP in allergic mouse mucus *in vivo*. **a.** MSD measurements from diffusion of 100 nm MIPs in tracheal mucus were used to calculate median pore sizes. **b.** Compared to healthy mice (grey, n = 7), mucus shifted towards tighter pore sizes with AOE exposure (cyan, n = 14). TCEP treatment (Tx), increased mucus pore sizes in AOE challenged mice (magenta) to be comparable to the healthy state. Values are means ± sem. P-values were determined by ANOVA with Tukey’s test for multiple comparisons.

**Suppl. Data Fig. 4.**
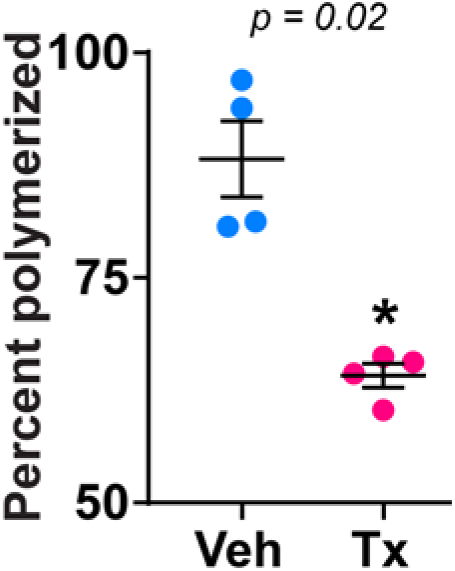
Relative quantification of mucin polymers from mice in AEC studies. Intensities of high molecular weight (polymerized) and low molecular weight (depolymerized) mucins were evaluated using Image Studio software (Li-Cor). Demarcations of polymerized and depolymerized signals are indicated in **Fig. 2c**. Data are presented as the fraction of polymerized versus total (polymerized and depolymerized) signals per lane. Data are means ± sem. Significance was determined by one-tailed Mann-Whitney U test with p-value shown and ‘*’ denoting significance (p < 0.05) from Veh.

**Suppl. Data Fig. 5.**
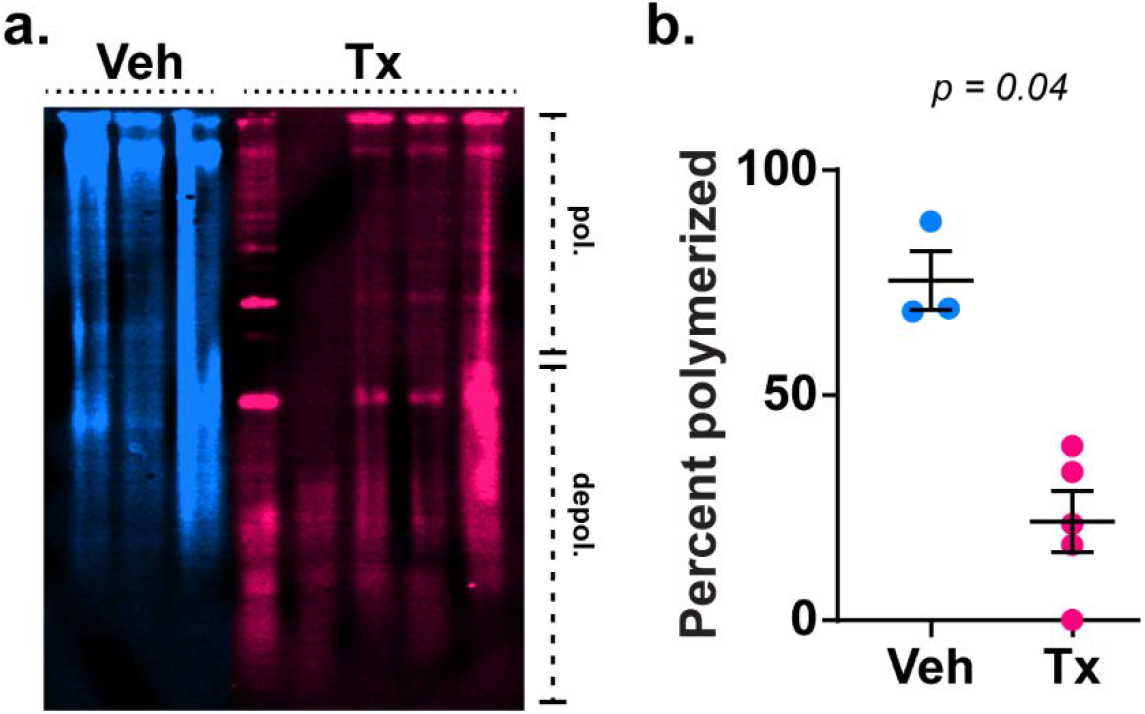
Immunoblot analysis of mucolytic effects in lung lavage fluid from allergic mice after AHR analysis. Lung lavage fluid from AOE challenged mice was obtained at the end of AHR studies. Equal volumes of lavage fluid (25 μl) were loaded per well in 1% SDS agarose gels. Samples were separated by electrophoresis, vacuum transferred to PVDF membranes, blotted with rabbit-anti-Muc5b, and detected with anti-rabbit IgG conjugated with Alexa 680. **a**. Relative to saline vehicle controls (Veh, cyan), Muc5b migration was enhanced by TCEP treatment (Tx, magenta). **b**. Intensities of high molecular weight (polymerized) and low molecular weight (depolymerized) mucins were evaluated using Image Studio software (Li-Cor). Demarcations of polymerized and depolymerized signals are indicated in **a**. Data are presented as the fraction of polymerized versus total (polymerized and depolymerized) signals. Data in **b** are means ± sem. Significance was determined by one-tailed Mann-Whitney U test with p-value shown and ‘*’ denoting significance (p < 0.05) from Veh.

**Suppl. Data Fig. 6.**
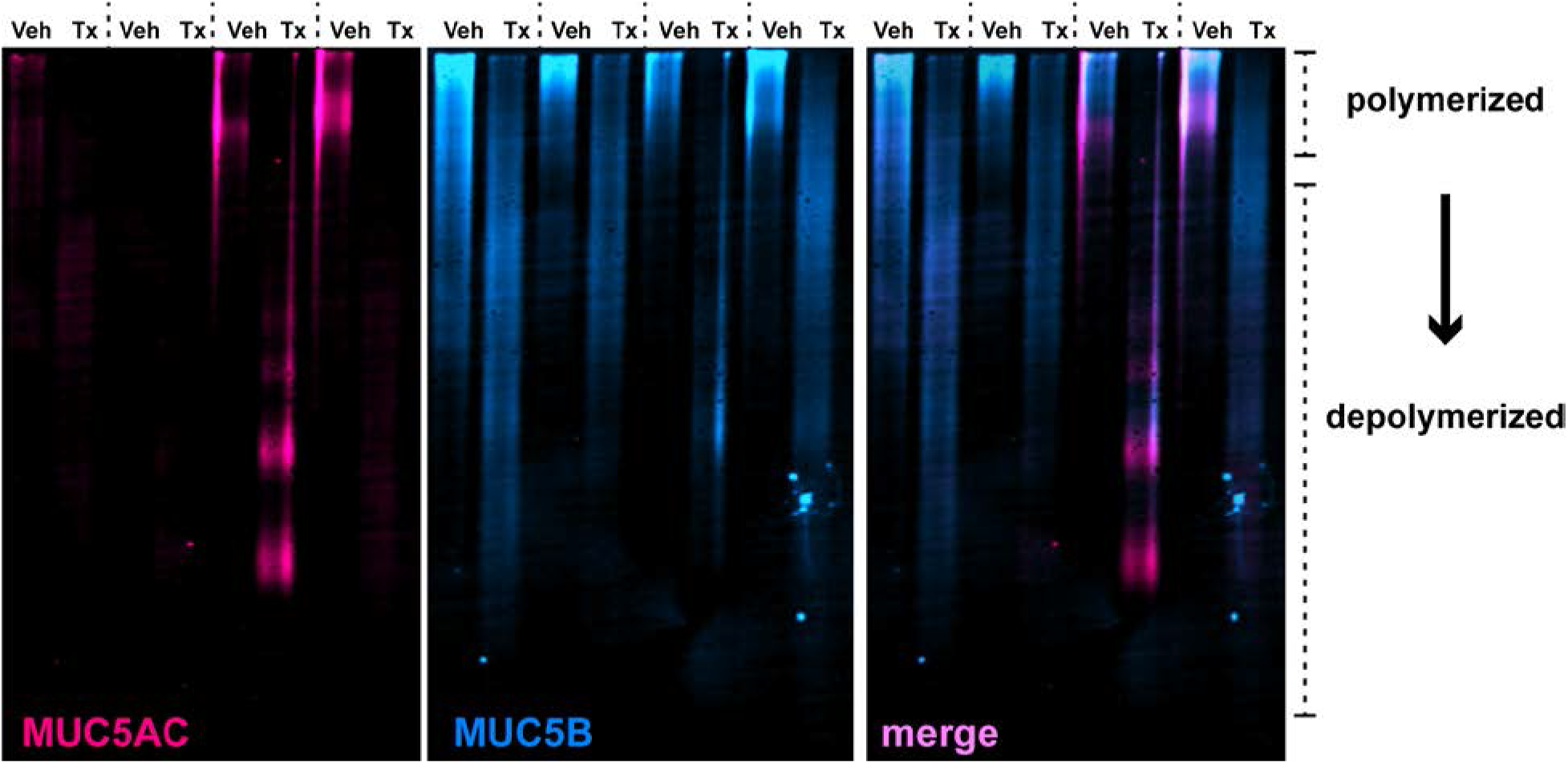
TCEP depolymerizes MUC5AC and MUC5B in bronchoalveolar lavage fluid from human asthma patients. Rabbit-anti-MUC5AC (magenta) and goat-anti-MUC5B (cyan) antibodies were applied (1:5,000 dilution) and developed using anti-rabbit-IgG conjugated with Alexa-680 and anti-goat-IgG conjugated with Alexa 800. Samples were treated with TCEP (10 mM final concentration) at neutral pH for 10 min at 37°C. Note that full reduction causes epitope loss for both anti-mucin antibodies, resulting in diminished signal intensities.

**Suppl. Data Fig. 7.**
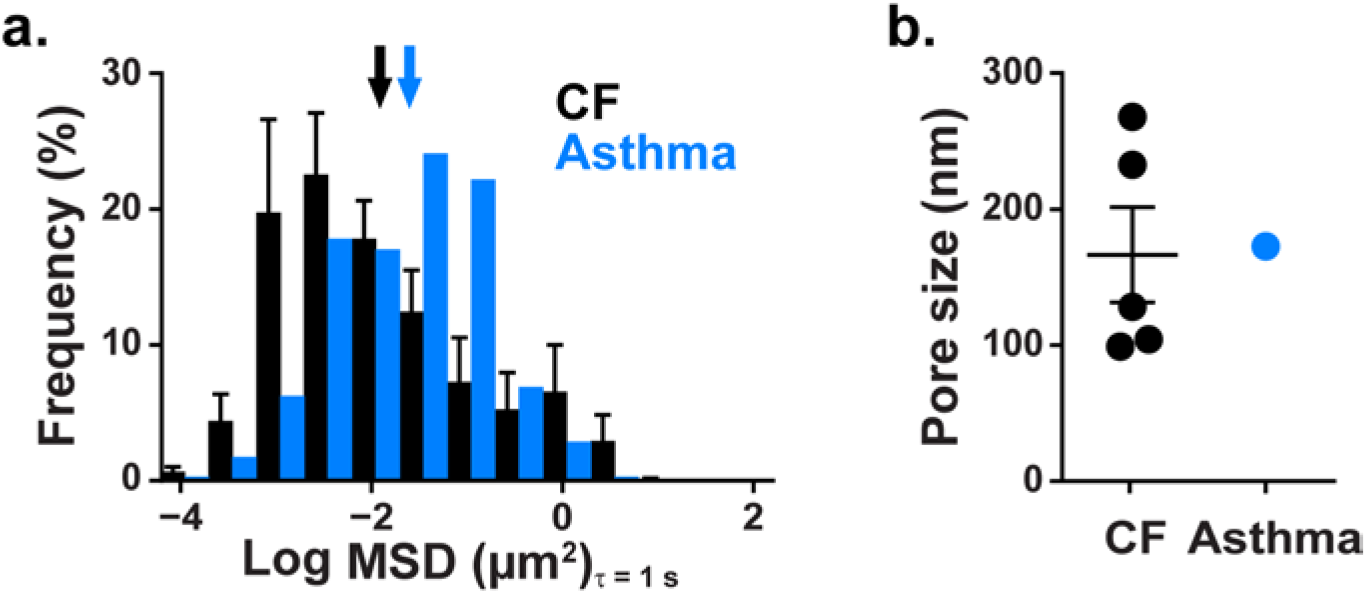
Histogram and medium pore sizes derived based from diffusion of 100 nm MIPs in spontaneously expectorated sputum from CF patients and in mucus aspirated from bronchial explants from a fatal asthma patient. CF sputum (n = 5), asthma (n = 1). **A**) Distribution of the log median MSD’s of individual MIPs. Data represent at least 500 particles tracked per sample. Inverted arrows indicate the average for each set. MSD values below Log (MSD) of −4 are below the tracking resolution. **B**) Median pore sizes in asthma mucus appear comparable to range in CF mucus. Data are means ± sem with dots showing individual sample medians in **b**. Mucus solids concentrations were 7.1 ± 0.5% (CF) and 7.7% (asthma).

**Suppl. Data Table 1.** Lavage leukocytes totals in AEC analysis.

|  | Measure | Vehicle, n = 6 | TCEP, n = 8 | p |
| --- | --- | --- | --- | --- |
| **Total cells** | mean (*sem*) | 15.6 (3.8) | 5.8 (0.8) | 0.03<sup>t</sup> |
|  | median (*quartiles*) | 13.7 (7.7-23.5) | 4.8 (4.2-6.9) | 0.006<sup>U</sup> |
| **Macrophages** | mean (*sem*) | 8.0 (1.8) | 4.2 (0.8) | 0.05<sup>t</sup> |
|  | median (*quartiles*) | 7.5 (4.9-10.7) | 3.5 (2.6-6.1) | 0.04<sup>U</sup> |
| **Eosinophils** | mean (*sem*) | 5.0 (0.2) | 1.0 (0.3) | 0.04<sup>t</sup> |
|  | median (*quartiles*) | 3.1 (2.1-8.2) | 0.6 (0.2-1.9) | 0.003<sup>U</sup> |
| **Lymphocytes** | mean (*sem*) | 1.4 (0.5) | 0.5 (0.08) | 0.07<sup>t</sup> |
|  | median (*quartiles*) | 0.8 (0.6-2.2) | 0.5 (0.3-0.7) | 0.02<sup>U</sup> |
| **Neutrophils** | mean (*sem*) | 1.2 (0.8) | 0.2 (0.08) | 0.1<sup>t</sup> |
|  | median (*quartiles*) | 0.4 (0.2-1.9) | 0.1 (0.05-0.3) | 0.05<sup>U</sup> |

<sup>t</sup> Welch's t test, one-tailed
<sup>U</sup> Mann-Whitney U-test, one-tailed

